# Electrokinetic active grooves for liposome capture, confinement, trajectory analysis and controlled release

**DOI:** 10.64898/2026.05.27.727973

**Authors:** Preethi Ravikumar, Matheus A. S. Pessôa, Sajad Shiekh, Zezhou Liu, Robert Sladek, Walter Reisner

## Abstract

Extracellular vesicles (EVs) are attractive candidates for minimally invasive disease monitoring. EV diagnostic signals often reside in low-abundant subpopulations, motivating single-vesicle measurements that can report distributions rather than ensemble averages. In practice, such assays rely on transient migration of a mobile vesicle through a detection region or performing cycled processing/measurement steps on vesicles permanently immobilized on surfaces via affinity-based capture. Single vesicle analysis would benefit from technologies that can combine characteristics of transient detection and surface immobilization to enable controlled capture, fixed time analysis and release of vesicles in a marker-neutral way. Here, we present electrokinetically active (RECON) grooves that provides low-voltage, programmable capture, timed retention, and release of nanoscale vesicles within microscopic grooves under continuous optical access. The platform uses a titanium embedded electrode with thin TiO_2_ passivation, enabling stable actuation over multi-hour experiments. Using fluorescent, size-defined liposomes as standards, we demonstrate (1) reversible, minute-long confinement at <5 V, (2) size-dependent confinement times, and (3) optical quantification of groove-guided drift velocities. Together, these results establish RECON grooves as a reusable, electrically programmable platform for controlled capture, timed retention, and release, while providing quantitative, trajectory-derived measures of confinement and transport.

## Introduction

Extracellular vesicles (EVs) are released continuously by cells and carry molecular cargo such as proteins, nucleic acids, and lipids, that can reflect the physiological state of their cell of origin.^1,2^ Because they circulate in biofluids, they are attractive targets for minimally invasive disease monitoring.^2,3^. EV associated molecular cargo provides a wide-range of biomarkers with diagnostic significance, particularly membrane based markers that are easily accessible and compatible with affinity probes such as antibodies and lectins. Affinity probes are used to capture, label, or classify vesicles based on membrane markers enriched in particular tissues or disease contexts.^2,3,4^. For example, in neurodegenerative diseases vesicles bearing CNS-associated surface features have been isolated from plasma.^5^ Similarly, in preeclampsia, placental-associated surface signatures are targeted to distinguish pregnancy-derived vesicles from the maternal background.^6^ In oncology and inflammatory disease, numerous assays likewise use binding to disease-linked antigens or glycan patterns displayed on the vesicle membrane.^3^

Most existing technologies with single EV analysis capability exploit either translocation through a detection region (transient-based, e.g flow cytometry) or permanent surface capture (immobilization based, e.g. ExoView).^3^ These two approaches entail a trade-off; transient approaches maximize throughput but minimize single vesicle interrogation times, while immobilization based approaches allow extended interrogation but sacrifice throughput and ability to perform downstream analysis/operations on captured vesicles. In contrast, trajectory-based measurements such as residence time, escape-time distributions, or position-dependent drift provide access to kinetics of confinement and transport within a programmable energy landscape, enabling measurements that are inaccessible to one-pass transit assays or permanently immobilized capture methods. For example, flow cytometry can deliver high-throughput multiparametric readouts,^7,8^ but each event is sampled only during brief passage through the interrogation volume; the measurement window is bounded by transit time, and the assay is inherently one-pass and thus does not support repeated measurements of individual vesicles.^7,8^ Other examples of transit-based technologies include pore-based resistive pulse sensing (RPS)^9^ and tunable RPS,^10^ but here too observation time is bounded by passage through the sensing region.^10–12^ Immobilization-based methods, such as single-particle interferometric reflectance imaging (SP-IRIS; ExoView-class assays)^13^ and related capture-array approaches,^13,14^ enable stable counting and phenotyping by immunocapturing vesicles onto a surface microarray.^15^ This approach can output multiple surface markers simultaneously and allows quantification of how many vesicles are positive for each marker.^13–15^ However, it imposes three constraints: (i) selection by capture chemistry (which biases what is observed), (ii) suppression of free motion (which precludes mobility-related measures), and (iii) retention without on-demand release (which prevents recovery of the assayed molecules).^13,15^ Controlled release following analysis could in principle enable repeated capture/analysis cycles to achieve high statistics and select sub-populations of interest.

Nanofluidic confinement is an attractive approach toward addressing these challenges, providing controlled vesicle handling and measurement under clinically relevant constraints: low-abundance and limited sample volumes.^3,16,17^ Nanofluidic devices impose steric confinement using pre-etched structures that confine molecules to lower-dimensional spaces (e.g., two-dimensional slits, one-dimensional nanochannels, and zero-dimensional cavities).^18–21^ These structures can hold individual vesicles in the field of view long enough to analyze their dynamics.^16,17^ However, most nanofluidic free-energy landscapes are determined by etched-in geometry and are therefore fixed once fabricated.^16,17^ This rigidity becomes especially problematic in EV workflows, where label multiplexing and improving signal-to-background often necessitates washing, labelling, or buffer exchange steps; such steps can modify EV surface state and ionic conditions, shifting electrokinetic forces and therefore the effective drive experienced by vesicles.^2,3,4^ These challenges motivate active confinement strategies in which the free-energy landscape can be tuned during the experiment instead of being built into the device.^22,23^ Active mechanical confinement can provide reversible trapping and release, but these approaches typically require extra actuation hardware, respond more slowly, create bulk flows that can deplete particle concentration, and are harder to standardize across devices.^23^

Active electrical control offers a complementary route to mechanical confinement. Because EVs (and vesicle standards) carry charge, their motion and confinement can be programmed using rapidly-tunable, externally-applied electric fields.^4^ For example, Anti-Brownian Electrokinetic (ABEL) trapping suppresses Brownian motion through real-time feedback control of an applied electric field and can hold a particle in view in free solution indefinitely.^24^ However, as the trapping mechanism in ABEL requires continuously adjusting dynamic transverse fields induced by external electrodes, this approach is difficult to adopt in an array format that could parallelize vesicle capture.

Here, we introduce electrokinetically active (i.e., RECON^25^) grooves for vesicle analysis. This approach implements programmable and reversible, low-voltage confinement and timed release. An enabling feature of our implementation is a titanium embedded electrode with thin TiO_2_ passivation. This supports low-voltage actuation in experiments spanning multiple hours with minimal degradation and enables repeated reuse of the same chip. To benchmark device performance, we performed experiments using synthetic fluorescent liposomes of pre-defined sizes.^2,11,15,26^ Our results demonstrate minute-long trapping at low voltages with simultaneous trajectory-based measurements that distinguish between sampled liposome sizes. Together, this work positions groove-based RECON lanes as a reusable, electrically programmable technology for vesicle manipulation with quantitative readouts, providing a foundation for capture → timed retention → release workflows applied to EV samples extracted from biofluids in liquid-biopsy based assays.

## Results

### Device design and fabrication

Devices (Fig. 1A) were fabricated on borosilicate glass by depositing Ti (80 nm) to form the bottom electrode, followed by SiNx (300 nm, PECVD) as an insulating layer. Photolithography and reactive ion etching (RIE) is used to pattern the SiNx into long, parallel grooves that expose the underlying Ti within each groove while leaving the surrounding surface electrically insulated. To suppress electrochemical degradation under applied fields, a thin TiO_2_ passivation layer (3 nm) was deposited by ALD.

**Figure 1.**
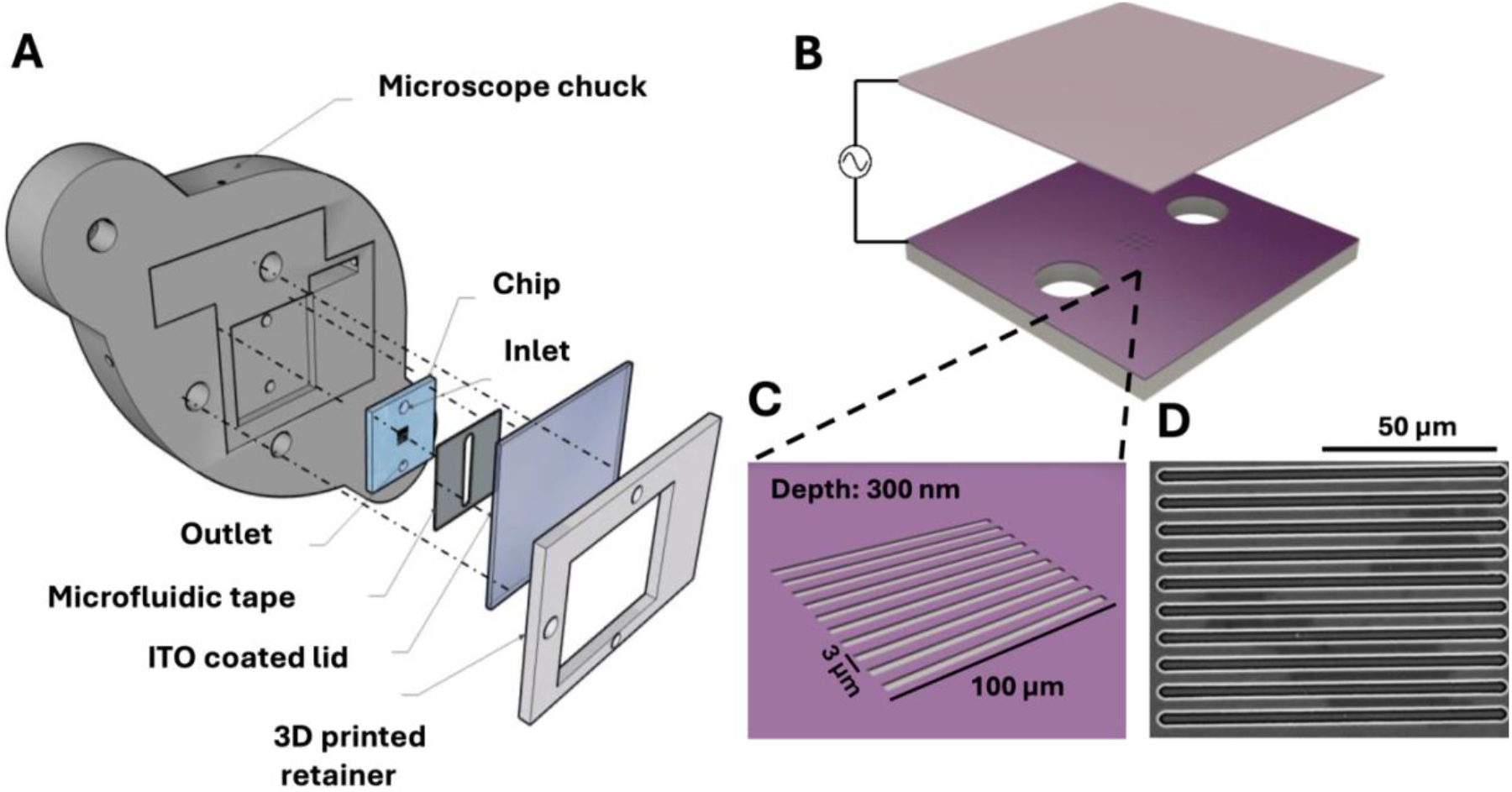
Device design and fabrication. (A) Schematic of the experimental assembly: the patterned chip is sealed with microfluidic tape and an ITO-coated lid to form a parallel-electrode flow cell with defined inlet/outlet fluidic access. The flow cell with embedded chip is then secured to a microscope chuck via a 3D-printed retainer. (B) Schematic of the electrode configuration used for groove-based reversible electrokinetic confinement (RECON): an AC potential is applied between a bottom electrode patterned beneath the grooves and an ITO-coated top lid, generating an electrically programmable confinement/transport landscape under optical readout. (C) Zoomed in view of the active region showing an array of long, parallel grooves that define the confinement lanes (w = 3 μm, h = 300 nm, L = 100 μm). (D) Representative SEM micrograph of the fabricated groove array illustrating the parallel-lane geometry used for vesicle capture, guided drift, and release.

The groove array defines the confinement geometry used throughout this work (w = 3 μm, h = 300 nm, L = 100 μm; Fig. 1B, C). For electrical actuation with optical readout, the patterned substrate was assembled into a parallel-electrode flow cell by bonding an Indium Tin Oxide (ITO)-coated coverslip as a transparent top electrode (Fig. 1B). Patterned adhesive tape both defined the microfluidic chamber for sample loading and exchange and set the electrode separation (d = 50 μm; Fig. 1A). Electrical connections to the Ti and ITO electrodes were made using silver epoxy (Fig. 1A).

Because trajectory-based metrics are interpretable only when the confinement geometry is known and consistent^16^, we verified groove morphology by scanning electron microscopy. SEM imaging confirmed long, parallel grooves with uniform width, spacing, and profile across the patterned region (Fig. 1D), supporting the treatment of each groove as a nominally equivalent lane for cross-site comparisons.

### Reversible single-vesicle confinement in grooves

A central goal of EV-oriented workflows is reversible single-molecule retention: vesicles must be held long enough for observation or perturbation but released without relying on surface binding^2,3^. We therefore tested whether our grooves support field-gated confinement at the single-vesicle level. We loaded the chamber with fluorescently labeled, negatively charged liposomes and applied an AC field across the parallel electrodes while imaging vesicle dynamics (Fig. 2A-D).

**Figure 2.**
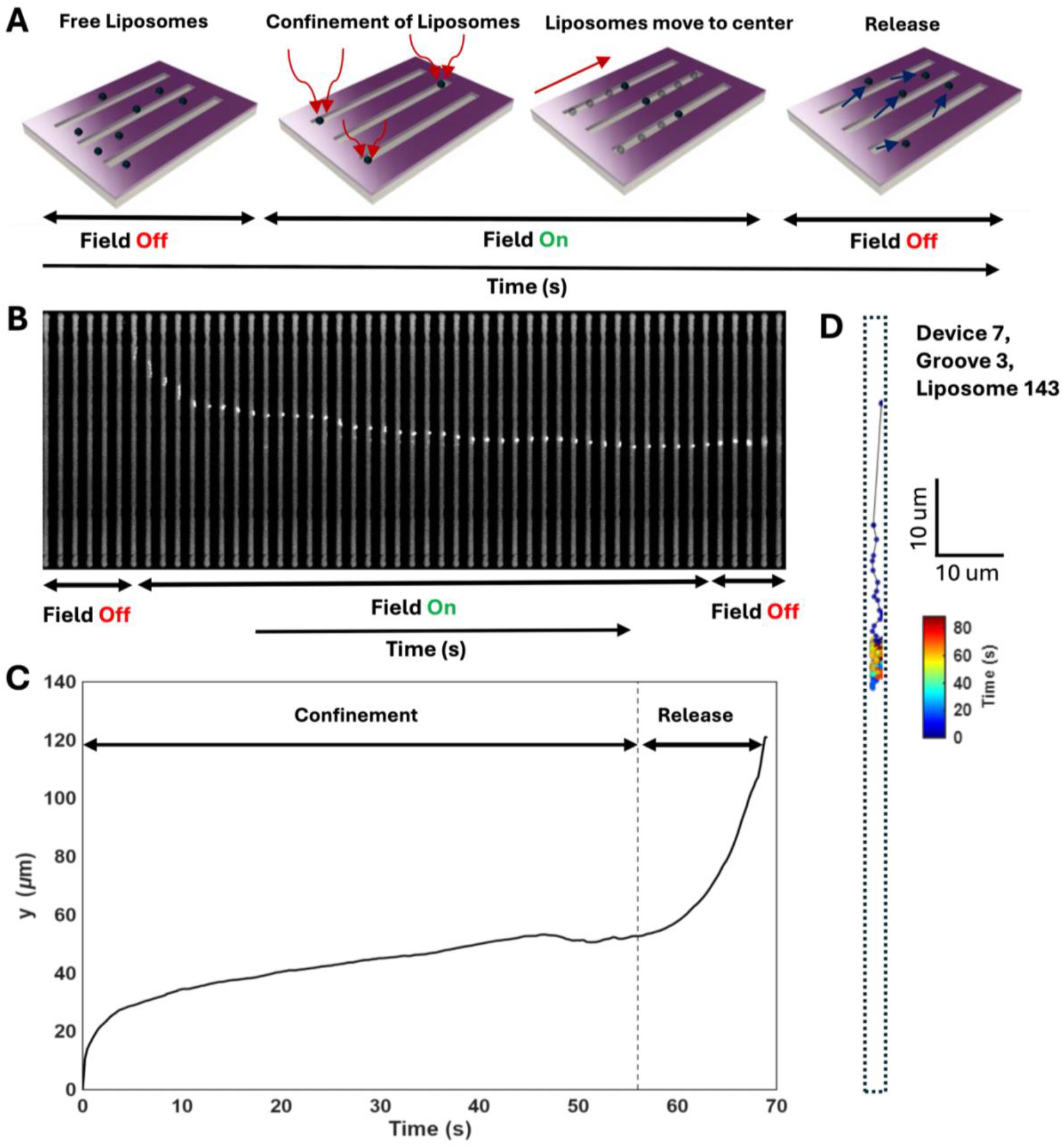
Reversible single-vesicle confinement in grooves at 10 kHz, 5 V, using 300 nm liposomes. **(A)** Cartoon showing device operating principle. With the field off, liposomes diffuse freely in bulk solution. When the AC field is applied, liposomes are captured into the grooves, undergo groove-guided drift toward the lane center, and then are released on demand by switching the field off. **(B)** Representative time-lapse kymograph (space–time montage) showing a single liposome trajectory across the groove array during a field-off → field-on → field-off sequence; confinement is maintained during the field-on interval and release occurs promptly after field deactivation. **(C)** Example of single-liposome axial position y(t) illustrating the confinement phase under field-on operation followed by rapid escape upon field removal (vertical dashed line marks the switching time). **(D)** Example single-vesicle trajectory within one groove (dashed outline), colored by time, demonstrating confined motion during the field-on window and the spatial localization achieved within the lane.

With the field off, liposomes underwent free diffusion without preferential localization in grooves (Fig. 2A). When the AC drive was applied (5 V), individual liposomes entered the grooves, remained laterally confined to a single groove, and progressed along the groove axis (Fig. 2B, D), yielding sustained single-vesicle trajectories (Fig. 2C). Turning the field off caused the same liposomes to exit the groove region and return to unconstrained diffusion (Fig. 2A-C). This reversible switching behavior is shown directly in Supplementary Video S1, where confinement and release are synchronized with the applied field state. Throughout the remainder of the paper, we use this reproducible switching to define a confined state (field ON) and a released state (field OFF), and we use it to generate time-resolved single-vesicle trajectories under controlled driving.

From these trajectories we extract three readouts: (i) a quantitative confinement metric based on escape-time under constant drive, (ii) a position-dependent axial drift profile along the grooves, and (iii) an effective electrophoretic mobility obtained under an experimentally calibrated electrical drive.

### Voltage-gated confinement at different voltages

To illustrate how electrical drive controls confinement at the single-vesicle level, we constructed time-lapse montages of an individual fluorescent liposome interacting with a groove under otherwise matched conditions, recorded over a fixed 90 s observation window at three applied voltages (5 V, 3 V, and 1 V; Fig. 3A-C). At 5 V, the liposome remained retained within the groove for a substantial fraction of the recording, yielding long uninterrupted in-groove intervals (Fig. 3A). Reducing the drive to 3 V shortened these in-groove residence intervals and increased the frequency of release events within the same 90 s window (Fig. 3B). At 1 V, retention was transient: the liposome only briefly entered the groove before exiting, producing short residence episodes and frequent loss from the groove (Fig. 3C).

**Figure 3.**
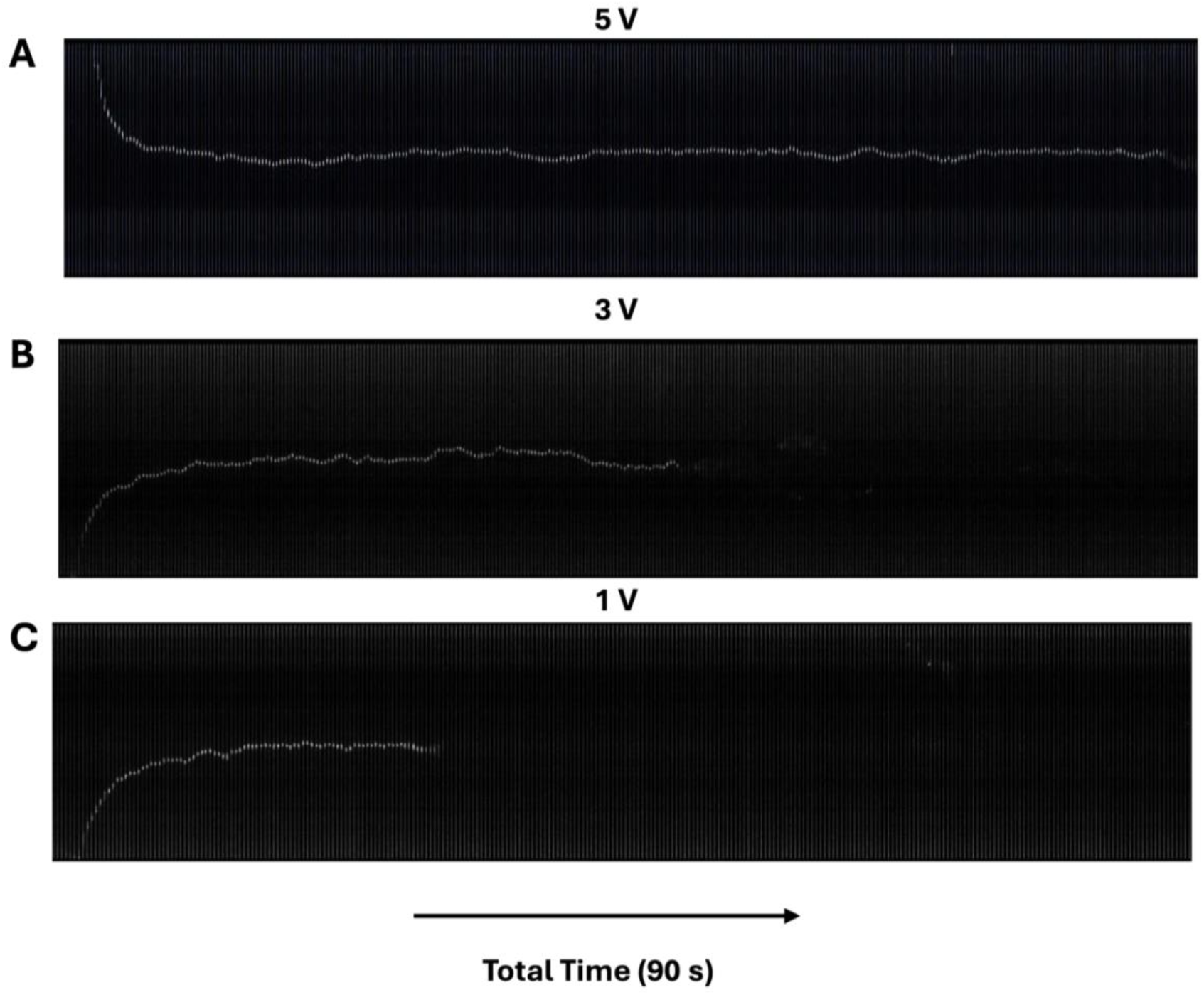
Voltage-tuned confinement dynamics in a single-liposome kymographs. Representative space–time montages (kymographs) showing single liposome trajectories in the groove array under different applied voltages over an identical acquisition window (total time 90 s). **(A)** 5 V, **(B)** 3 V, and **(C)** 1 V. Increasing the applied voltage strengthens the electrokinetic confinement, yielding longer retention within the lane and more sustained groove-guided motion during the observation window, while weaker drive produces earlier loss/escape from the confined region.

Together, these montages provide a direct visual demonstration that groove confinement is electrically gated and tunable, with residence time decreasing systematically as the applied voltage is reduced (5 V > 3 V > 1 V) over the same acquisition duration (Fig. 3A-C).

### Quantifying confinement with escape-time measurements

To quantify confinement, we defined an event-based escape time t_esc_, giving the time interval between vesicle entry into the groove and vesicle exit while the drive remains on (Fig. 4A). For each voltage and vesicle size, we obtained the distribution of t_esc_ from single-vesicle trajectories (Fig. 4B). Across conditions, escape times were well described by an exponential distribution, consistent with a homogenous Poisson escape process. We therefore fit a single-exponential model

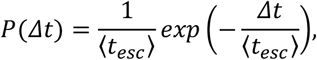

and report the mean escape time ⟨t_esc_⟩ (Fig.4B). We found that confinement strengthened systematically with electrical drive for all three vesicle sizes, but the magnitude of the effect depended strongly on size (Fig. 4B). The smallest vesicles (100 nm) showed a modest increase in retention as voltage was raised (⟨t_esc_⟩ = 8.7 s at 1 V to 16.4 s at 5 V; ∼1.9×), whereas the larger vesicles responded much more sharply to the same increase in drive: 200 nm vesicles increased from 12.7 s to 58.3 s (∼4.6×), and 300 nm vesicles increased from 15.3 s to 73.2 s (∼4.8×). At any fixed voltage, escape times also followed a consistent size ordering (300 nm > 200 nm > 100 nm) (Fig. 4B). For example, at 5 V, 300 nm vesicles remained confined for 73.2 s on average versus 58.3 s for 200 nm and 16.4 s for 100 nm; i.e., the largest vesicles were retained ∼4.5× longer than the smallest under the same applied drive, highlighting a systematic size dependence of retention and release kinetics.

**Figure 4.**
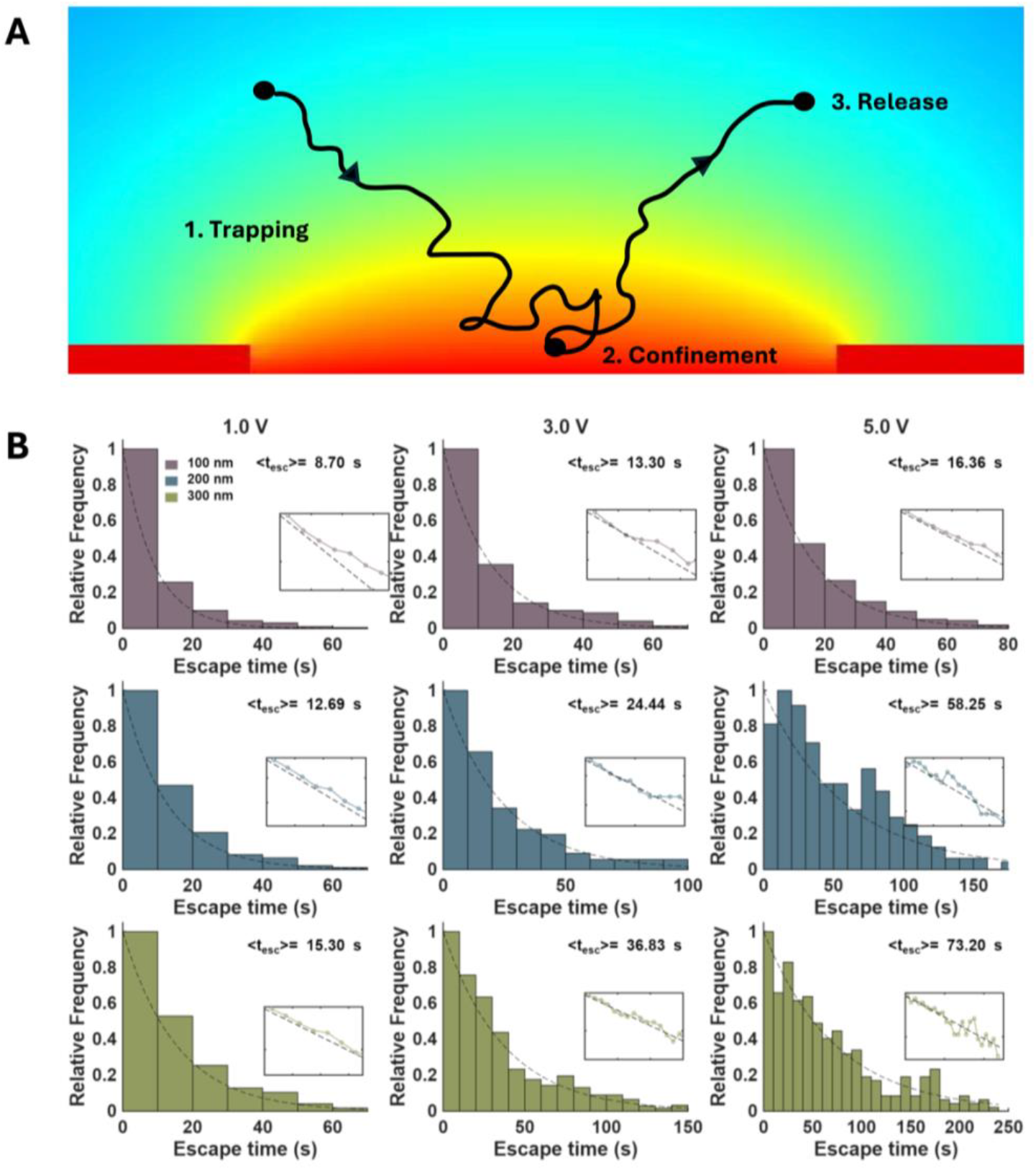
Quantifying confinement strength using escape time measurements. **(A)** Conceptual trajectory illustrating the event definition used for escape-time analysis: a vesicle is captured into the groove (trapping), remains confined under the applied field (confinement), and subsequently exits the groove while the drive remains on (release/escape). **(B)** Escape-time distributions for three liposome sizes (100, 200, and 300 nm; rows) measured at three applied voltages (1, 3, and 5 V; columns). Histograms show relative frequency of escape times *t*_esc_; dashed curves indicate single-exponential fits consistent with an approximately constant escape rate, and the reported ⟨*t*_esc_⟩ values summarize the mean escape time for each condition. Insets show the corresponding log-linear representation used to assess exponential behavior.

### Confinement scales with device-measured current

Because conductive buffers and electrode impedance make nominal voltage an imperfect proxy for the local electrical drive, we parameterized each condition by the device-measured current^25^. A finite time-averaged current through the assembled device necessarily implies a finite time-averaged current through each electrokinetically active groove, and therefore a non-zero current density *J* on the groove floor. In a simplified electrostatic picture, this can be implemented as a Neumann boundary condition that specifies the normal electric field at the groove floor in terms of the local current density, *E*_*n*_ ≈ *J*/*σ*, where *σ* is the solution conductivity^25^. The relevant vesicle dynamics occur on ∼ 0.1s (or longer) time scales, far slower than the 10 kHz actuation period (100 μs), so from the perspective of capture and confinement the system responds to the time-averaged electrical drive. For each applied drive, we recorded an I–V curve on the same chip and extracted the corresponding current *I* (Supplementary Fig. S1), which serves as a direct readout of the electrical response under the measurement conditions.

When plotted against current density, the mean escape time ⟨*t*_esc_⟩ increases monotonically for all three vesicle sizes (Fig. 5). This preserves the same size ordering observed under voltage control (300 nm >200 nm >100 nm) at comparable current, while making clear that the stronger applied drive systematically increases confinement. Notably, the separation between size classes is maintained across the full current range: at a given current density, larger vesicles exhibit longer residence times, indicating robust, size-dependent retention and release kinetics in the electrically programmed landscape (Fig. 5). By relating confinement to a measured electrical quantity rather than nominal bias, this representation supports quantitative comparison across chips and days in which voltage-to-field transfer may vary.

**Figure 5.**
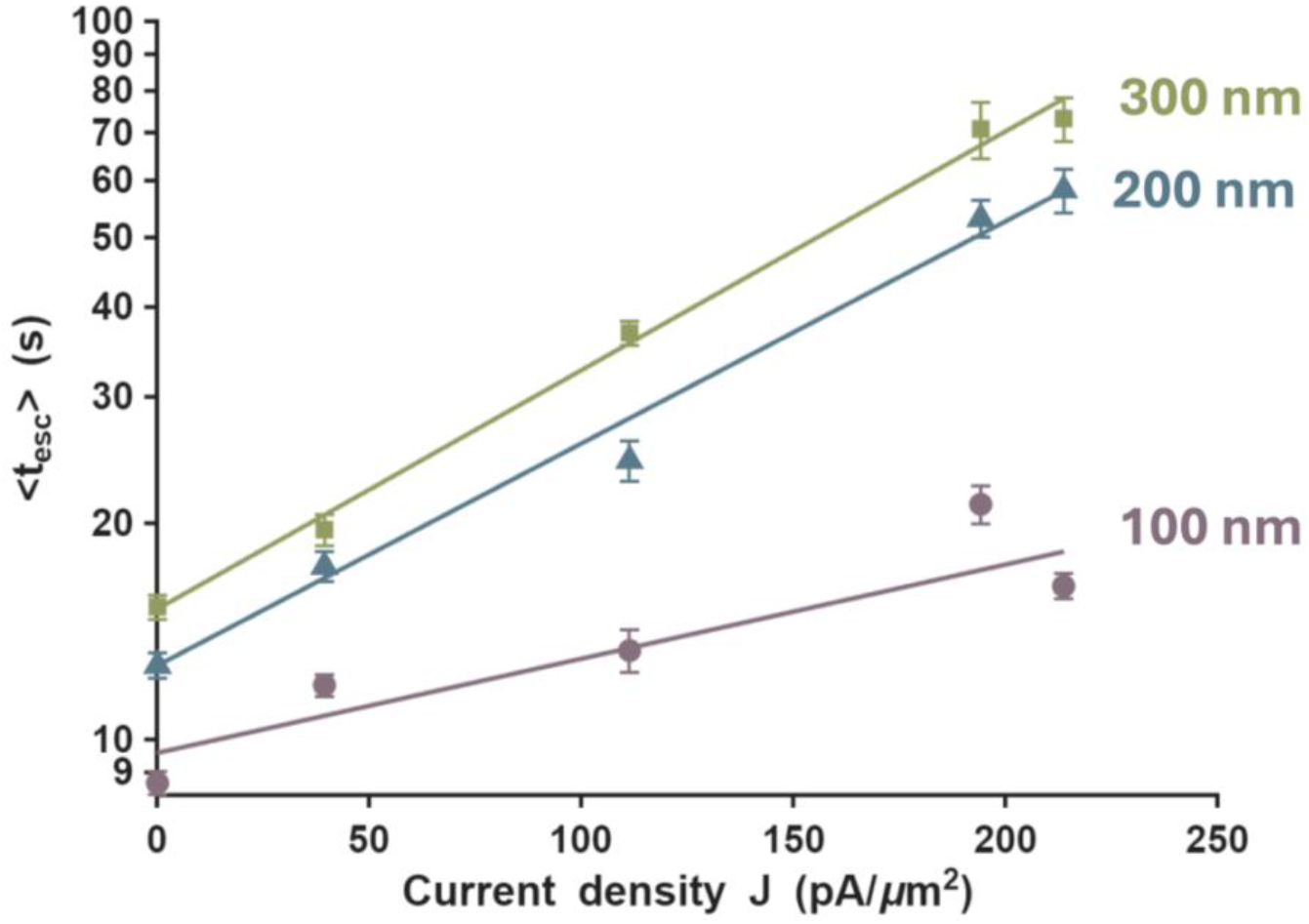
Confinement scales with electrical drive and depends strongly on vesicle size. Mean escape time ⟨*t*_esc_⟩ plotted as a function of measured device current density *J* for three liposome sizes (100, 200, and 300 nm). Points show ⟨*t*_esc_⟩ extracted from single-exponential fits to the escape-time distributions (Fig. 4) with error bars indicating uncertainty in the fitted mean for each condition. Lines are linear guides/fits highlighting the monotonic strengthening of confinement with increasing electrical drive and the pronounced size dependence, with larger liposomes exhibiting substantially longer mean residence times at the same current density.

The slopes in Fig. 5 provide additional insight into how the electrical landscape strengthens with applied drive. Because the escape-time distributions are well described by a single-exponential, escape is consistent with a Poisson process characterized by a rate *k*, with ⟨*t*_*esc*_ ⟩ ≈ 1/*k*. In this regime, confinement can be interpreted as a barrier-controlled escape process in which the escape rate depends exponentially on an effective barrier height Δ*U*:

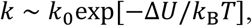

so that

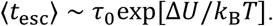

Within this framework, the rise of ⟨t_esc_⟩ with current indicates that applied drive increases the effective barrier; equivalently, the apparent slope of ⟨t_esc_⟩ versus *I* reports how rapidly the trapping barrier deepens with the imposed drive.

Interpreting *I* as a proxy for the time-averaged electrical drive (and thus for a characteristic field scale through *J* ∼ *I*/*A* and *J* = *σE*), the size dependence of the slopes indicates size-dependent sensitivity of confinement to applied drive. Larger vesicles show a stronger increase in ⟨*t*_esc_⟩ with *I*, consistent with an effective barrier Δ*U*(*I*) that grows more rapidly with drive for larger particles.

### Position-dependent drift velocities along grooves and relation to simulated field profile

To relate groove-guided drift to the spatial structure of the electrical drive, and to define the velocity input used for the mobility extraction below, we quantified the mean axial drift speed as a function of position along the groove, *v*(*y*), under a fixed actuation condition (5 V, 10 kHz) (Fig. 6A). We plot one half of the groove (entrance → midpoint), where *y* denotes distance from the channel midline along the groove axis. All three vesicle sizes exhibit the same strongly structured drift landscape: motion is fastest near the entrance and decreases monotonically as vesicles approach the midpoint, tending toward near-zero drift at the center (Fig. 6A). This consistent slow-down across 100, 200, and 300 nm indicates that vesicles experience a position-dependent drive along the groove rather than a spatially uniform electrophoretic drift.

**Figure 6.**
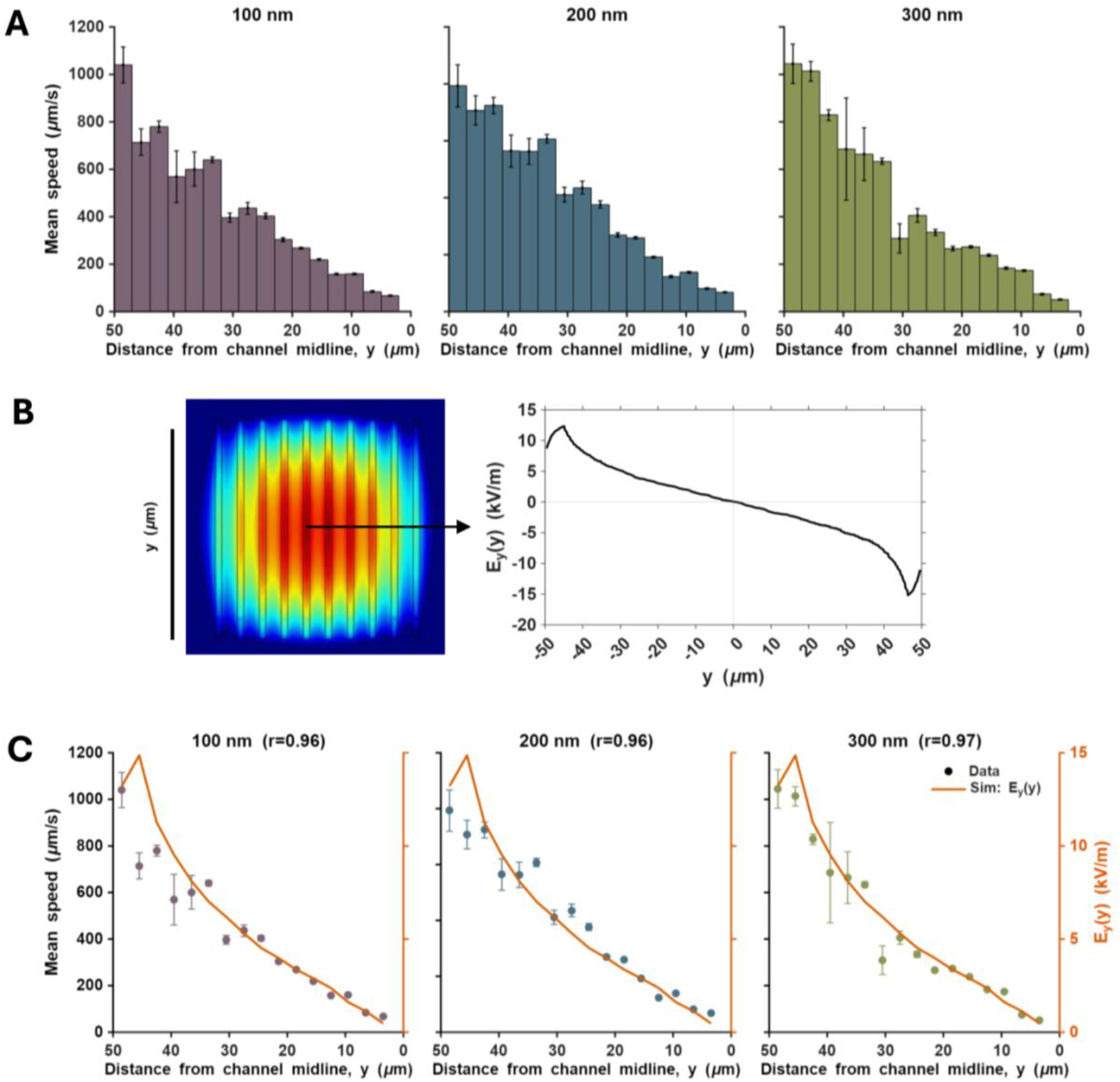
Position dependent groove-guided drift velocities and relation to simulated field profile. (**A)** Mean axial drift speed as a function of position along the groove, plotted versus distance from the channel midline, *y*, for three liposome sizes (100, 200, and 300 nm; left to right) under a fixed actuation condition (5 V, 10 kHz). Bars show binned mean speeds; error bars indicate standard error within each position bin. Across sizes, drift is fastest near the groove entrance (large ∣ *y* ∣) and decreases toward the midline, approaching near-zero speed at the channel centre. **(B)** COMSOL simulation of the groove-region electric field (left) and the extracted axial field component *E*_*y*_(*y*) along the drift direction (right), shown in IV-calibrated units (kV/m). **(C)** Measured mean drift speeds (points with error bars; left axis) overlaid with the IV-calibrated simulated field profile (solid curve; right axis) for each liposome size, demonstrating that the spatial decay of drift speed mirrors the position-dependent electrical drive; panel titles report the correlation coefficient *r*.

We next tested whether this position dependence is set by the electrical landscape imposed by the groove geometry. We modelled the field profile in COMSOL (Electric Currents, frequency domain, 10 kHz) under an Ohmic bulk-conduction assumption (Fig. 6B). To avoid introducing an explicit assumption about interfacial voltage drop at the Ti surface, the simulation was implemented with a boundary condition of fixed current density: a prescribed normal current density was applied on the exposed Ti region, the ITO electrode was grounded, and remaining boundaries were insulating. The resulting simulated *E*_*y*_(*y*) was scaled using the experimentally determined, effective electrical drive in the device. Specifically, the current density from the IV measurements together with the independently measured buffer conductivity *σ*_meas_ = 0.0351 S/m define an effective bulk field *E*_eff_ = *J*/*σ*_meas_, which we use to report the simulated field profile in IV-calibrated units (Fig. 6B,C).

The IV-calibrated simulation predicts an axial driving field whose magnitude progressively decreases toward an electric potential maximum at the groove midpoint (Fig. 6B). Note that for the negatively charged liposomes examined here, electrophoretic drift is opposite the electric-field direction. A decreasing ∣ *E*_*y*_(*y*) ∣ predicts a corresponding reduction in drift speed as the midpoint is approached. Consistent with this expectation, the measured drift profiles closely track the IV-calibrated field magnitude along the groove (Fig. 6C), with strong correlations for each size class (Pearson *r* = 0.96 for 100 and 200 nm; *r* = 0.97 for 300 nm). The simulated profile also shows a small increase in field magnitude near the groove edges that is not clearly resolved in the experimental drift curves. The uncertainty in the measured velocities is largest near the groove edges, consistent with the larger instantaneous velocities expected in these regions under the stronger local field. This increases variability in the measured drift speeds which might be preventing us from resolving the small near-edge increase predicted by the simulation. Nevertheless, the overall agreement in these results shows that the dominant, systematic variation in groove-guided drift speed is governed by the electrical landscape predicted by the groove geometry and provides the IV-calibrated field input required for the device-calibrated mobility analysis that follows.

### Effective electrophoretic mobility in grooves using an experimentally calibrated field drive

Finally, we asked whether groove-guided drift can provide an electrophoretic mobility readout in the same lane used for confinement. We estimated an effective on-chip mobility by normalizing the measured drift to the experimentally calibrated field scale^27^,

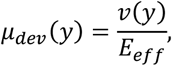

where v(y) is the mean axial drift velocity at position y along the groove and E_eff_ is the current-calibrated electrical drive defined for each condition. The electrical drive is obtained by recording an I–V curve on the same chip used in experiments, computing the current density J = I/A_eff_, and then relating the current density to electric field using an independently measured bulk conductivity *σ*_bulk_ ^28^:

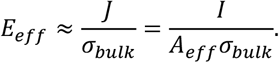

The resulting mobilities fall in the expected 10^−9^ m^2^ V^−1^ s^−1^ range and show a modest decrease in magnitude with increasing vesicle size (Fig. 7). Using the current-calibrated field scale, the on-chip mobilities were *µ*_dev_ ≈ 32.3 ± 2.1 × 10^−9^(100 nm), 31.0 ± 2.2 × 10^−9^(200 nm), and 28.8 ± 2.7 × 10^−9^ m^2^ V^−1^ s^−1^(300 nm) (mean ±SEM; Fig. 7). Independent bulk electrophoresis measurements (Zetasizer) yielded mobilities of comparable magnitude, *µ*_bulk_ = (38.5 ± 2.2) × 10^−9^ (100 nm), (37.0 ± 2.2) × 10^−9^ (200 nm), and (33.3 ± 2.2) × 10^−9^ m^2^ V^−1^ s^−1^ (300 nm) (Fig. 7). Comparing methods reveals a consistent, size-dependent offset: *µ*_dev_ is lower than *µ*_bulk_by 6.2 × 10^−9^ (16%) at 100 nm, 6.0 × 10^−9^ (16%) at 200 nm, and 4.5 × 10^−9^ m^2^ V^−1^ s^−1^ (14%) at 300 nm (Fig. 7). Overall, current-calibrated groove drift provides a device-integrated mobility readout in the correct physical range with a reproducible size dependence, while remaining broadly consistent in magnitude with bulk electrophoresis.

**Figure 7.**
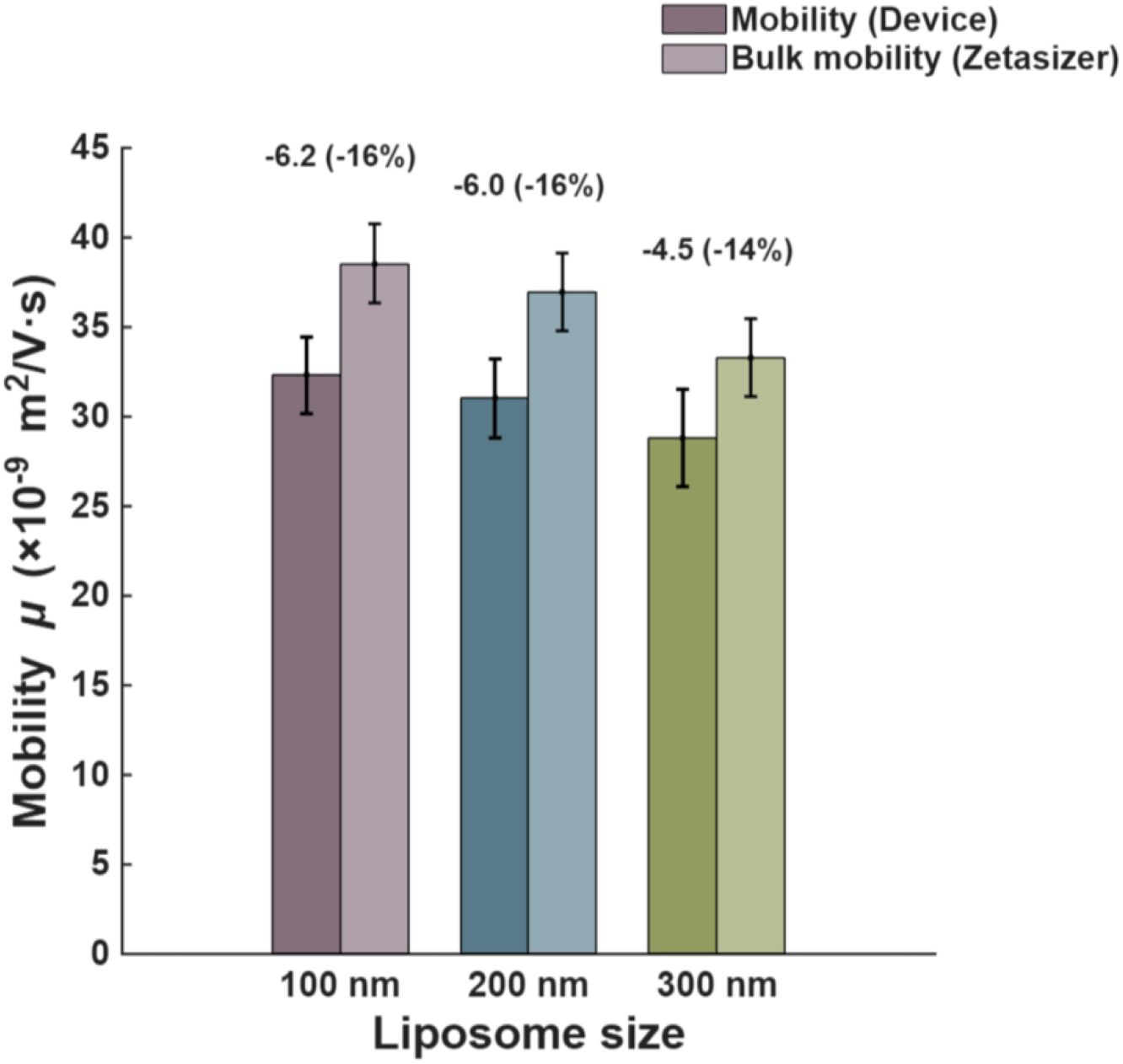
Device-calibrated mobility from groove-guided drift agrees with bulk electrophoresis. Effective electrophoretic mobility *µ* extracted from groove-guided drift in the RECON lane (Mobility (Device)) compared with independent bulk mobility measured by a Zetasizer for three liposome sizes (100, 200, and 300 nm). Bars show mean mobility values with error bars indicating measurement uncertainty. Numbers above each size indicate the absolute and percent difference between the device-derived and bulk values, demonstrating close agreement across sizes and supporting groove-guided drift as a quantitative mobility readout that can be compared directly with bulk electrophoresis under calibrated electrical drive.

Notably, our device measured a modest size-dependent mobility, consistent with the Zetasizer measurements. This modest size dependence is physically plausible at our ionic strength. At *I* ≈ 1mM, the Debye length in water is 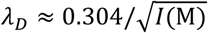, giving *λ*_*D*_ = 9.6 nm and therefore *κa* = *a*/*λ*_*D*_ ≈ 5.2, 10.4, 15.6 for 100/200/300 nm vesicle diameters (radii *a* = 50, 100, 150nm)^29^. Under these finite-*κa* conditions, classical electrokinetic theory relates electrophoretic mobility to both zeta potential, *ζ*, and the Henry function *f*(*κa*) (i.e., *µ* ∝ *ζ f*(*κa*)), with the size-independent Smoluchowski limit recovered only at *κa* ≫ 1 ^30,31^. However, at fixed *ζ*, the Henry-function correction alone would predict only a modest size dependence and does not by itself explain the observed decrease in mobility with increasing vesicle size. The measured trend therefore suggests that, in addition to finite-*κa* effects, other factors such as size-dependent differences in effective *ζ* or device-specific interfacial effects may contribute.

In addition, for vesicles the effective *ζ* inferred from electrophoresis is known to be sensitive to the measurement medium and interfacial chemistry (including bound ions and near-surface effects), and EV/liposome *ζ* is commonly reported to depend on ionic strength and sample handling/preparation ^4,32^. Taken together, the shared size trend between *µ*_dev_ and *µ*_bulk_ (Fig. 7), combined with the finite-*κa* regime at 1 mM, supports treating the observed decrease in *µ* with size as a reproducible electrokinetic signature under our buffer conditions.

## Discussion

In this work, we demonstrate that groove-based RECON can both control and quantify nanoscale vesicles under electrical actuation. By applying the electric field within a defined geometry, the RECON grooves enable reversible capture of individual liposomes with tunable, minute-long confinement and trajectory-based quantification. This electrically programmable device thus has the potential to measure and hold vesicles in view during EV workflow steps such as solution exchange or reagent exposure, without relying on permanent immobilization.^3,4^

Overall, our results highlight two core strengths of RECON: (i) programmable confinement as an assay-handling operation, and (ii) electrical calibration that provides reliable comparison across experiments. First, RECON grooves achieve reversible confinement across multiple vesicle sizes (Fig. 4, 5). This establishes the lane as a programmable state sequence: capture → timed retention → release, implemented within a defined geometry. For EV workflows, this enables repeated cycling (capture/hold/release) without committing vesicles to permanent surface binding, and could allow vesicles to be held through deliberate perturbations prior to release for downstream processing.^3,4^ Second, to make confinement and drift metrics more comparable across chips and days, our RECON strategy utilizes drive calibration, which relies on both individual chips’ I–V relationships and independently measured buffer conductivities. This approach mitigates the ambiguity in the effective electric field and thus enables quantitative comparisons of confinement strength that would otherwise be obscured when plotting solely versus nominal bias. Taken together, these two features position RECON grooves as a promising basis for clinically relevant EV workflows.^2,3^

Beyond assay handling, these RECON grooves also have the potential for simultaneous, quantitative measurement, yielding trajectory-derived readouts (e.g., residence/escape measurements and groove-guided drift). Simultaneous optical measurements across particles allows us to construct escape-time distributions, a capability with clinical relevance because EV subpopulations are often expected to appear as subtle shifts in distribution within heterogeneous samples.^3,33^ If these subpopulations possess distinct surface properties, those differences could alter electrokinetic confinement behavior and thereby produce corresponding shifts or secondary modes in the escape-time distributions measured here. Consistent with this, community guidance (ISEV/MISEV) emphasizes EV heterogeneity and the need for measurements that report distributions in a way that remains comparable across workflows.^2^ In future work, it will therefore be important to test whether the retention and escape-time distributions accessible in RECON lanes can be used to resolve subpopulations in mixed EV samples.^2,3^

Having established that the grooves can report trajectory-based metrics, a natural next step for quantitative electrokinetic studies in micro- and nanofluidic systems is to map those observables directly onto the calibrated local field landscape.^28^ Here, we found that the measured position-dependent velocity profiles were consistent with the spatial structure predicted by our simulations of an electrophoretic model.^30,31^ This suggests that electrophoresis captures a dominant component of groove-guided drift under our conditions.^30,31^ However, we do not necessarily infer a single mechanism from this agreement. Prior work from our lab on electrokinetic confinement in nanocavities argued that confinement was electrophoretic, but also noted that electroosmosis, while unlikely to drive capture from bulk, could still contribute secondary, geometry-specific flows that influence in-trap distributions.^25^ Related devices have also leveraged polarization forces directly: DEP has been used as an explicit operating principle for nanoscale confinement/manipulation in related micro/nanofluidic geometries (e.g., frequency-programmed DEP confinement in the 10–100 kHz range).^34^ In the groove-lane geometry here, analogous EO/DEP contributions could in principle modify trajectories without necessarily disrupting the overall velocity– position trend. Disentangling the residual contributions of other forces, through targeted frequency sweeps, surface-coating tests to modulate electroosmosis, and comparisons to extended models incorporating EO/DEP terms remains an important direction for future work.

With respect to the device’s design, several extensions could be pursued. One is lane-level addressing, for example by segmenting the embedded electrode into independently driven lanes or serial segments along a single lane. This would enable on-chip parallel multi-lane operation, with adjacent lanes running different retention programs or waveform protocols on the same input sample, allowing internal controls and direct, side-by-side comparisons of how retention kinetics depend on drive or buffer conditions without separate devices or runs. A second extension is to operate grooves as programmable transport lanes. Because vesicles already exhibit groove-guided drift, the lane could be used as a conveyor for staged processing—for example capture near an entrance, translation to a defined interrogation zone, a timed hold for imaging, and then release. With segmentation, sequences such as capture → move → hold → move → release become possible, creating a device-level analogue of a timed incubation line in which vesicles can be exposed to reagents, washed while retained, and then transported to a downstream region for readout or collection.

Overall, groove-based RECON lanes provide a novel electrically programmable platform for controlled capture, timed retention, and release of nanoscale vesicles under optical access, while yielding calibrated, trajectory-based metrics of confinement and transport.

## Materials and Methods

### Device design and fabrication

Devices were fabricated on borosilicate glass substrates. A titanium bottom electrode (80 nm) was deposited by e-beam evaporation. A silicon nitride insulating layer (SiN_x_, 300 nm) was deposited by plasma-enhanced chemical vapour deposition (PECVD). Long, parallel grooves were patterned into the SiN_x_ by photolithography followed by reactive-ion etching, exposing the underlying Ti only within each groove while leaving the surrounding surface electrically insulated. Groove dimensions were, *w* = 3 μm *h* = 300 nm, and *L* = 100 μm. To suppress electrochemical degradation under applied fields, an thin TiO_2_ passivation layer (3 nm) was deposited by atomic layer deposition (ALD).

### Flow-cell assembly and electrical connections

For optical readout under electrical actuation, the patterned substrate was assembled into a parallel-electrode flow cell using an indium tin oxide (ITO)-coated coverslip as the transparent top electrode. Patterned adhesive tape defined the chamber footprint and set the electrode separation to *d* = 50 μm. Electrical contact to the Ti and ITO electrodes was made using silver epoxy. The assembled device was driven by a function generator (Agilent 33220A), and the applied waveform and device response were monitored using a digital oscilloscope (Rigol DS1074Z).

### Liposome preparation and fluorescent labelling

Synthetic fluorescent liposomes were used as EV-sized standards. 100 nm and 200 nm DOPC/CHOL liposomes labelled with Rhodamine-DHPE were purchased from FormuMax (catalog nos. F60103F-R and F60103F2-R, respectively) and used as received. Stock suspensions were diluted into the measurement buffer immediately prior to experiments and handled according to the manufacturer’s storage recommendations. Also, 300 nm liposomes were prepared by membrane extrusion using DOPC/Cholesterol (70/30 mol%) with Rhodamine-DHPE (0.5 mol%) for fluorescence imaging. To confer a net negative surface charge, we incorporated a carboxylate (–COOH)–functional lipid (deprotonated to –COO^−^ under experimental conditions), ensuring an electrophoretic response under the applied electric fields. Liposomes were stored at 4°C. Prior to loading, liposome suspensions were diluted to 50 nM concentration in measurement buffer.

### Buffers, conductivity measurement, and device loading

Experiments were performed in 1 mM TBE (pH 8). The bulk conductivity, *σ*_bulk_, of the measurement buffer was measured immediately prior to experiments using an Oakton pH/Conductivity Meter (510 series). Devices were primed with buffer to remove air and equilibrate surfaces, then loaded with liposome suspension by pipetting into the inlet reservoirs.

### Electrical actuation protocol

An AC drive was applied across the parallel electrodes at 10 kHz using a sinusoidal waveform with a 0 to +V amplitude (unipolar offset; i.e., the voltage oscillated between 0 and +V). All voltage values denote this 0→+V peak voltage. Field-gated confinement was defined operationally by switching the drive on (confined state) or off (released state) during imaging. For confinement-versus-drive experiments, the voltage was stepped from 1 to 5 V (0→+1 V, 0→+2 V, …, 0→+5 V) under otherwise fixed buffer and imaging conditions.

### Fluorescence microscopy and imaging conditions

Liposome dynamics were imaged on an inverted fluorescence microscope (Nikon Eclipse Ti). Fluorescence excitation and emission were selected to match the lipophilic dye (Rhodamine-DHPE) using an appropriate rhodamine/TRITC filter set at 575 nm wavelength. Time-lapse image sequences were acquired at a 30 ms frame interval for durations sufficient to capture multiple confinement and escape events per condition. Experiments were performed at room temperature.

### Single-vesicle detection and trajectory extraction

Single-vesicle trajectories were obtained from fluorescence time-lapse image sequences. Vesicles were detected as diffraction-limited spots and linked across frames using custom MATLAB/ DoM pipeline. Each trajectory was assigned to a groove lane by spatial localization relative to the groove array orientation; trajectories outside grooves were treated as free diffusion controls when needed.

### Escape-time definition and fitting

To quantify confinement, an event-based escape time *t*_*esc*_ was defined as the duration between a vesicle entering a groove and subsequently exiting while the drive remained on. Entry and exit were detected from the trajectory’s lateral position relative to the groove boundaries within groove ROI. For each vesicle size and drive condition, distributions of t_esc_were compiled across events. Escape-time histograms were fit to a single-exponential model

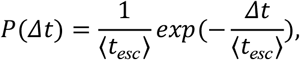

and the mean escape time ⟨*t*_es*c*_ ⟩ was reported for each condition. Fits were performed by maximum likelihood and confidence intervals were estimated by bootstrap.

### Device electrical response and current-based parameterization

For each chip, the electrical response was measured under the same buffer conditions used for trapping experiments by recording an I–V curve across the electrode pair. For each applied amplitude, the corresponding device current I was extracted and used as an experimental descriptor of applied drive for that condition. Where required for field scaling, the effective current density was computed *as J* = *I*/*A*_*eff*_, with *A*_eff_ defined as the exposed electrode area participating in conduction. The parameter *A*_eff_ was obtained from the CAD pattern used to produce the photolithography mask and fabricated features. Bulk conductivity *σ*_bulk_ was measured independently, and an effective field scale was defined by *E*_eff_ ≈ *J*/*σ*_bulk_.

### Drift velocity profiles along grooves

To characterize groove-guided transport, axial drift velocity v(y) was computed as a function of position along the groove axis. Trajectories were rotated into the groove coordinate system and binned by axial position y using bins 3 µm in width. Within each bin, the mean axial velocity was computed from frame-to-frame displacements divided by the frame interval. Velocity profiles were computed for each vesicle size under fixed actuation (5 V, 10 kHz) and plotted over one half-groove length from entrance to midpoint, exploiting symmetry about the groove center where appropriate.

### COMSOL simulation of the electrical landscape

The spatial structure of the driving field was modelled in COMSOL Multiphysics using the Electric Currents interface in the frequency domain at 10 kHz under an Ohmic bulk-conduction approximation. The ITO top electrode was grounded, and all non-electrode boundaries were set to electrical insulation. The exposed Ti region within the groove was assigned a current-driven boundary condition (Neumann condition) specifying the normal current density. The resulting potential and electric-field distributions were computed and the axial field component along the groove was extracted for comparison to measured *v*(*y*). The device current (or current density) measured under the experimental buffer conditions was used as an experimental proxy for applied drive. Combined with the independently measured buffer conductivity, this defined an effective field scale via *E*_eff_ = *J*/*σ*, where *J* is the device-measured current density and *σ* is the buffer conductivity. The simulated field profile was then IV-calibrated so that its magnitude over the experimental region of interest was consistent with this effective field scale, yielding a calibrated field profile for plotting and quantitative comparison. Agreement between measured drift profiles and simulated field shape was quantified by linear regression between *v*(*y*) and the simulated field magnitude, reporting Pearson correlation r.

### Effective electrophoretic mobility from groove-guided drift

An effective on-chip electrophoretic mobility profile was computed as

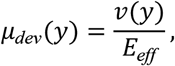

where *v*(*y*) is the mean axial drift velocity at position *y* and *E*_eff_ is the current- and conductivity-based field scale defined for the corresponding condition. For each vesicle size, a single summary mobility *µ*_*dev*_was reported as the mean of *µ*_*dev*_(*y*) over a defined analysis region (entrance-to-midpoint), with uncertainty reported as the standard error of the mean (SEM) across trajectories.

### Bulk electrophoresis measurements

Independent bulk electrophoretic mobility measurements *µ*_bulk_were obtained using a Zetasizer Ultra in the same buffer used for on-chip experiments. Liposome suspensions were diluted to the manufacturer-recommended concentration range and measured in the capillary cell at room temperature. Mobilities were reported as mean ± SEM.

### Statistics and Data analysis

The experimental videos were preprocessed using Fiji ImageJ macros for region of interest selection, noise subtraction, and frame averaging. The postprocessing was performed in MATLAB, through a customized data analysis pipeline for particle tracking, plotting, fitting, and processing.

## Acknowledgements

Funding is provided by Natural Science and Engineering Research Council of Canada (NSERC, RGPIN-2025-06363 and ALLRP 597782-24), the New Frontiers for Research Fund-exploration (NFRFE-2023-00526) and the Fonds de recherche du Québec-Nature et technologies (FRQ-NT, 329544). The authors also acknowledge the Nanotools-Microfab at McGill University for use of their fabrication resources and the facility for electron microscopy research (FEMR) for use of electron microscopes.

## Supplemental Figure

**Supplementary Figure S1.**
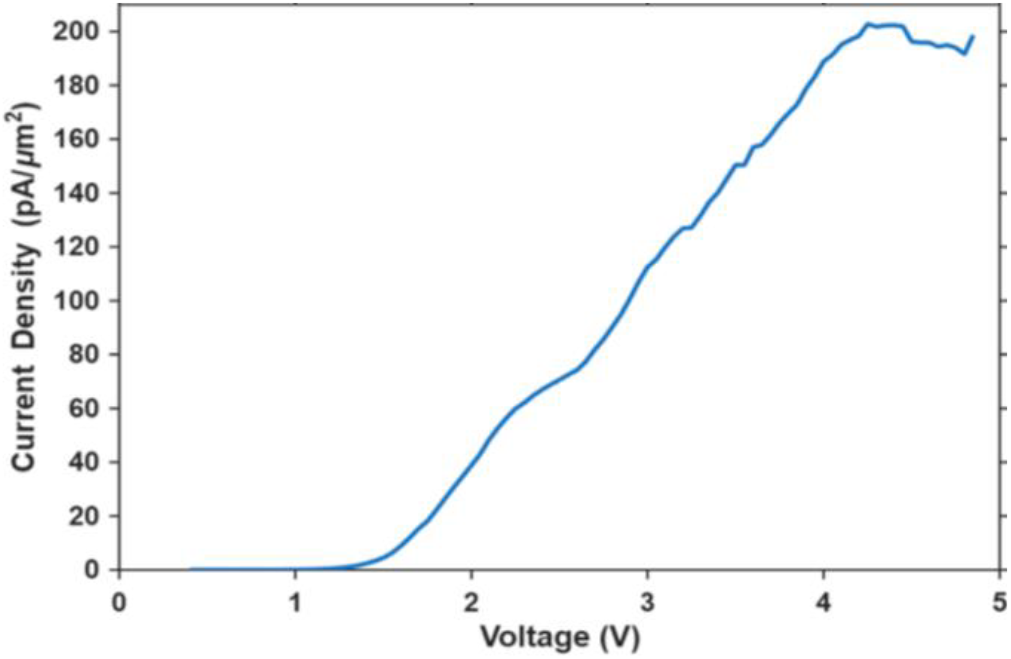
Device electrical characterization (I-V curve). Current density measured through the grooves as a function of applied voltage under the experimental buffer conditions (1 mM TBE, pH 8, bulk conductivity *σ* = 0.0351 S/m, room temperature), using a AC drive at 10 kHz with electrode separation d = 50 μm and groove geometry w = 3 μm, h = 300 nm, L = 100 μm. The nonlinear increase in current density with voltage reflects the combined effects of buffer conductivity and the device/electrode impedance and motivates using the device-measured current (or current density) as an experimental proxy for the effective electrical drive in subsequent analyses.

## Supplemental Video

**Supplementary Video S1**. Reversible electrokinetic confinement and release of 300 nm fluorescent liposomes within an electrokinetically active groove at 10 kHz and 5 V in 1 mM TBE. The video shows groove-guided liposome motion before field application (“Field OFF”), confinement during electrical actuation (“Field ON”), and release following field removal (“Field OFF”). Field-state labels are overlaid directly on the video frames.

